# Development of a new militarily-relevant whole-body low-intensity blast model for mild and subconcussive traumatic brain injury: Examination of acute neurological and multi-organ pathological outcomes

**DOI:** 10.1101/2021.09.15.460417

**Authors:** Sarah C. Hellewell, Ibolja Cernak

## Abstract

This work describes a newly developed experimental mouse model reproducing features of blast-induced neurotrauma (BINT), induced in operationally relevant manner using a compressed air-driven shock tube. Mild BINT (smBINT) was induced by one exposure to a low-intensity blast (LIB), whereas subconcussive BINT (rscBINT) was caused by repeated exposures to LIB.

To mimic an operational scenario when a soldier is standing when exposed to blast using a quadruped experimental animal (mouse), a whole-body holder was developed to position mice in a bipedal stance, face-on toward the pressure wave generated in a shock tube. This restraint avoids ‘bobble head’ movement, thus prevents tertiary blast effects, and allows administration of fast-acting inhaled anesthetics *via* nose cone.

Using this model, we established and validated paradigms for primary blast-induced mild and repetitive traumatic brain injuries Our results showed that a single exposure to 69 kPa (10 psi) was capable of inducing smBINT, whereas three-rounds of exposure to 41 kPa (6 psi) caused rscBINT.

Mice recovered rapidly from both types of BINT without prolonged neurological dysfunction. Mild superficial pathology was found predominantly in the lungs 24h after injury, with equivalent pathology after smBINT or repetitive rscBINT. The Purkinje layer of the cerebellum exhibited neuronal damage persisting up to 7d. Similar to some other models as well as clinical findings, this model reproduces blast-induced cerebellar pathology. In conclusion, this model positioning mice in a bipedal stance and facing front-on toward the shockwave provides realistic representation of operational scenarios and reproduces militarily-relevant smBINT and rscBINT in the laboratory.

## Introduction

Blast-induced neurotrauma (BINT) can occur during military training or in theatre, as well as a result of mining, chemical or other civilian incidents^1-4^. Blast-induced neuropathology was not always considered a unique physical entity^5,6^, although the term ‘shell shock’ has been used for decades to describe the behavioural and neurological consequences of blast injury in the absence of obvious trauma^7,8^.

Clinically, BINT manifests in prolonged deficits in the neurological, cognitive and psychological spheres^9,10^, with patients reporting loss of balance and coordination alongside deficits in executive function, memory, learning, attention, and emotional processing, as well as mood disorder and social isolation^11-14^. BINT is now recognised as a complex multifactorial injury^15^, with clinical and preclinical research defining varied pathological consequences, including disruption of the blood-brain barrier^16,17^, edema^18^, hypometabolism^11^, apoptosis^19,20^, axonal dysfunction and white matter abnormalities^21-23^, glial activation^24^, and generation of pathological forms of tau^23,25^, among others^25,26^. Many reports have now established BINT pathology as particularly prominent in the cerebellum^11,27,28^, suggesting that there may be structurally-specific pathological consequences.

Mechanistically, BINT may be due to passage of the blast wave through the skull and tissue displacement via acceleration and/or rotation of the head, or by transfer of kinetic energy from the blast wave through large blood vessels in the abdomen and chest to the central nervous system (CNS)^15,29^. This hydraulic interaction initiates oscillating waves that traverse the body at approximately the speed of sound in water, delivering the kinetic energy of the blast wave to the brain^30^ and causing morphological and functional damage to distinct brain structures^5^. These methods of blast injury are not mutually exclusive^3^ and are likely synergistic; indeed, preclinical data implicate both the direct interaction of the wave with the head^23^, and shockwave-induced vascular load ^31^ in the pathogenesis of BINT.

While the importance of multi-system, multi-organ response to blast exposure is more obvious in moderate-to-severe brain injuries, it is often neglected in the case of mild BINT. However, accumulating evidence demonstrates that systemic alterations initiated by blast significantly influence the brain’s response, contributing to the pathobiology of acute and/or chronic deficits due to blast^32^. It has been hypothesized that hyperinflation of the lungs caused by blast overpressure stimulates the juxtacapillary J-receptors located on the alveolar interstitial surface and innervated by vagal fibers, triggering a vasovagal reflex that leads to apnea followed by tachypnea, bradycardia, and hypotension^32-35^. In addition, hypoxia/ischemia caused by the pulmonary vagal reflex stimulates chemoreceptors in the left ventricle, activating a cardiovascular decompressor Bezold–Jarish reflex leading to significant parasympathetic efferent discharge to the heart^36^. This may cause further bradycardia and peripheral blood vessel dilatation, eventually contributing to cerebral hypoxemia.

Several preclinical models of BINT have been developed in rodents^18,28,37-39^ and larger quadrupedal animals^40-42^. A commonality of blast modeling to date has been a naturalistic positioning of the animal in a quadrupedal stance, with the head as the first bodily structure to interact with the blast wave. While neuropathology is observed in these models in a generally graded response to injury severity, body positioning has been shown to have vastly different transfer of dynamic pressure whether the animal is facing head front-on, side-on or away from the blast^43^. There have been few models in which animals are positioned in a bipedal stance, in which the blast wave would interact with the head and body simultaneously in a more realistic interpretation of human body position^44,45^. Therefore, the goal of this study was to develop an operationally relevant mouse model of BINT in which the mouse is placed upright to best reproduce the consequences of the blast wave as it would interact with the body in an upright position in man. As most BINT injuries are mild^39^, we examined the acute physiological, systemic and neuropathological response in paradigms of single mild BINT (smBINT) and repetitive subconcussive BINT (rscBINT).

## Methods

### Construction of a mouse holder for whole body blast exposure

The mouse holder was based on a currently implemented design used in the study of blast effects on cultured brain cell aggregates^46^. This initial structure was chosen as it provided minimal resistance to the blast wave during propagation along the blast tube. Aluminum arms connected by a central anchor enable the holder to slide inside the blast tube (Figure 1).

**Figure 1.**
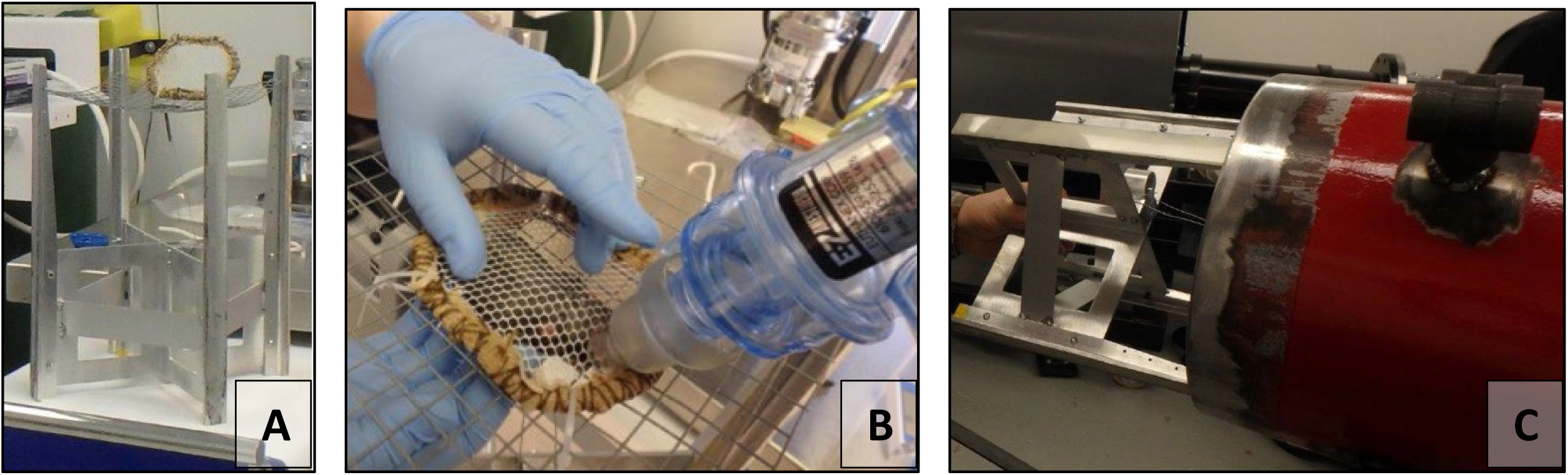
Mouse holder and restraint. The structure (A) comprised four rigid arms to which a sheet of Hardware Wire was secured to support the animal in a bipedal position inside the ABS. (B) a ring of fabric mesh with a flexible barrier adjustable to body size holds the animal firmly without placing undue pressure on limbs. This covering also allowed controlled ventilation via nose cone. Prior to BINT, the holder was tilted horizontally so the mouse was facing towards the blast wave in a bipedal stance (C) and advanced to 30 cm from the end of the tube.

### Selection and appraisal of restraint materials

The optimal material was identified as a strong loose-weave netting which could restrain the entire body in a single sheet and would not impede passage of the blast. The netting was sewn around a Gear Tie (strong, flexible rubber coated wire) fashioned into an oval shape and reinforced with a strong crepe material and thick cotton stitches (Figure 1). The netting cover was attached at four points with cable ties, with two in permanent placement to act as a hinge, allowing the covering to be placed precisely over the mouse. This restraint apparatus provided support and stability for the mouse during the blast, with high-speed footage revealing minimal movement of the head, limbs or torso during the blast wave. An additional benefit of this apparatus was the suitability for continuous anaesthesia administration via nose cone during securement and final adjustments prior to blast.

### Determination of blast parameters

The Advanced Blast Simulator (ABS) at DRDC Suffield allows generation of precise blast waves of known intensity corresponding to injuries on a spectrum from mild to severe. The 24-foot ABS generates a shockwave using helium compressed to a pre-determined threshold in a driver (compression) chamber, which, when the critical pressure threshold is reached, causes the rupture of polyethylene sheets placed between the driver and the driven (wave propagation) sections of the ABS. The polyethylene sheets are used in predetermined combinations, with the pressure at rupture dependent on thickness of the polyethylene in a linear fashion. In contrast to other shock tubes, the DRDC Suffield ABS contains both a hex-divergent driver to better model the spherically symmetric characteristics of a blast wave, as well as a rollaway anechoic end-wave eliminator to prevent reflection of the blast wave. The ABS is also equipped with five side-mounted pressure sensors along the length of the driven section to record the static pressure and wave propagation, as well as viewing ports to enable high-speed videography. The DRDC Suffield ABS was previously used to model various intensity blasts in rats, however no parameters of mild and subconcussive blast had been defined for mice. To examine appropriate driver pressure, various combinations of 0.003” and 0.005” polyethylene sheets were assessed in combination with various driver pressure thresholds, until the correct combinations were found to reliably reproduce waves corresponding to 6, 8, 10 and 15 psi, based in literature representations of mild BINT. The Canadian Army and US Army Doctrines specify exposure limits of 3 and 4 psi (respectively) for overpressure during range training^47,48^, however helmet-mounted sensors have detected frequent repetitive exposures at or above 6 psi^49-51^. There is now evidence that repetitive subconcussive blast events may elicit the same symptoms as those seen for mTBI, particularly in the operational scenario in which repeated blasts are experienced within a short timeframe. Our parameters for mild and subconcussive injury were guided by the literature describing other shock tube systems, for which the overpressure range is posited to be between 5-11 psi^15^, and mimics the operational range of 7.5-15 psi^52^. Preliminary blast experiments were performed to determine appropriate loading conditions in which mild and subconcussive effects might be modeled. For the purpose of these experiments, we defined ‘subconcussive’ injury as the highest exposure threshold (kPa) as which a single exposure produced no observable acute neurological effects nor systemic pathological consequences (examined using the Yelverton blast injury scoring system, described below). Mild injury was defined as the highest kPa at which mild systemic pathology could be observed. 6 and 10 psi met the criteria for subconcussive and mild intensity blasts, respectively, with 8 psi also approximating mild injury, and 15 psi causing moderate systemic pathology.

### Subconcussive blast intensity parameters

To achieve a reproducible repetitive subconcussive intensity blast (rscBINT), 6 psi (41 kPa) was set as the blast intensity. Three 0.003” membranes were used to create a diaphragm of 0.009” total thickness. To reach the appropriate driver pressure, the threshold was set to 42.9 psi (296 kPa), with an average time to rupture of 14.5 milliseconds.

### Mild intensity blast parameters

To model single-blast induced mild BINT (smBINT), 10 psi (69 kPa) was set as the blast intensity. Three sheets of 0.005” thickness and one 0.003” sheet were used to create a diaphragm of 0.018” total thickness. The driver pressure required to rupture this membrane was found to be 78 psi (538 kPa), with an average time to rupture of 15.06 milliseconds.

### Animal experiments

All mouse experiments were conducted in accordance with Canadian Council on Animal Care (CCAC) guidelines, and were approved by both the University of Alberta and Defence Research and Development Canada (DRDC) Suffield Animal Ethics Committees. Adult male C57Bl/6 mice (22 ± 2g; Charles River Laboratories, Montreal, Quebec, Canada, n=64) were housed in ventilated, temperature-controlled rooms on a fixed 12-hour light/dark cycle with access to food and water *ad libitum*. Prior to injury induction, mice were randomly allocated to either a) single 10 psi blast exposure (n=24), b) three 6 psi exposures (n=24) or c) sham control (n=16). Mice designated for repetitive blast exposures were anaesthetised and prepared as described below, with a second exposure two hours after their initial blast injury, and third 24 hours after the first blast.

### Blast procedure

Mice were anaesthetised with 3% isoflurane evaporated in 30% oxygen & 70% nitrous oxide in an induction chamber for two minutes, followed by an additional four minutes of administration via nose cone while the animal was secured to the blast holder. When appropriately secured and the mouse deeply anesthetized (confirmed by the absence of tail, corneal and toe pinch reflexes), the mouse holder was dispatched to the ABS and rotated so the mouse was in a bipedal position, facing front-on to the blast wave. In this stance, the holder was rapidly placed into the ABS and advanced 30 cm from the end of the tube (2.4 metres from the membrane) (Figure 1). The end-wave eliminator was quickly secured, and the blast sequence initiated. After blast exposure or sham conditions, mice were rapidly removed from the holder by cutting the cable ties on one side and placed into a clear Perspex cage for acute neurological monitoring.

### Acute neurological recovery

Immediately after removal from the holder, mice were continuously assessed for recovery of tail, corneal and toe pinch reflexes, as well righting reflex. Recovery monitoring also included time to groom and time to display exploratory behaviour. Mice were also assessed for changes in body weight prior to and after a single blast exposure as well as between rscBINT exposures.

### Perfusion and Tissue Preparation

At 1 day and 7 days mice were deeply anaesthetised with sodium pentobarbital and placed in an empty cage until loss of pedal and corneal reflexes. Transcardial perfusion was performed with 10 ml of ice-cold saline followed 8 ml of 10% formalin. Brains were rapidly removed and placed in 10% formalin for a further 12-16 hours post-fixation before immersion in 30% sucrose for 3-4 days for cryoprotection prior to freezing. Following this, brains were washed in phosphate buffered saline (PBS), frozen on dry ice and stored at −80°C until sectioning. The cerebellum of each mouse was sectioned at 30μm and stored in 12-well culture plates in a cryopreservation buffer until use.

### Yelverton blast pathology scoring

The gross pathological consequences of whole-body blast exposure were identified after perfusion using the Yelverton blast scoring system ^45,53^. This system provides an excellent profile of whole-body damage by combining scores for the extent (E) of organ damage, the grade (G) of organ damage, the severity (ST) of pathology, and the severity depth (SD) to indicate the depth of disruption of the worst pathology observed. These scores are combined in the following formula to reach a final injury severity (IS) score: IS = (E + G + ST) x SD.

### Haematoxylin and Eosin stain

Cerebellar sections were stained with Haematoxylin and Eosin (H&E) to examine gross morphological changes. Sections were mounted to Superfrost Plus slides (Fisher Scientific, Ottawa, Ontario, Canada) and air dried. Once dry, sections were rehydrated in running tap water, before 3-minute changes in increasing concentrations of ethanol (70, 100%). Next, sections were incubated in Haematoxylin stain for 3 minutes, rinsed in running tap water until the water ran clear, destained in acid alcohol, and placed in Eosin stain for 5 minutes, then dehydrated in xylene prior to coverslipping.

### Statistical Methods

Statistical analysis of all data was performed using GraphPad Prism version 6.0 for Macintosh (GraphPad, La Jolla, CA). All data was assessed by 2-way ANOVA, designed to determine differences between level of blast exposure (sham, rscBINT, smBINT) and time after injury. Where overall significance was detected, post hoc Tukey or Dunnet’s t-tests were employed to determine significance at individual treatment levels between timepoints.

## Results

### Blast wave characterization

Blast wave parameters were established for 10 and 6 psi exposures as representatives of mild intensity single blast and subconcussive repeated blasts, respectively. Polyethylene plastic membranes in predetermined combinations were used to generate the peak overpressure, with three sheets of 0.005” thickness and one sheet of 0.003” thickness (combined thickness 0.018”) found to generate a blast wave of 10 psi (69 kPa). To reach this pressure, the driver pressure threshold was set at 78 psi (538 kPa), with an average time to rupture 0f 15.06 milliseconds. The pressure was monitored at several gauges along the tube (Figure 2A), with a gauge directly in line with the mouse when placed in the holder to determine the pressure at that level of the blast tube. The dashed line indicates the moment of membrane rupture, at which the pressure in the driver section rapidly decreases while the pressure in the driven section increases along the blast tube. The data generated from sensor ‘ts4280’ at the level of the mouse was further mapped as a pressure trace to determine the blast wave characteristics of a 6 psi (Figure 2B) and 10 psi (Figure 2C) wave.

**Figure 2.**
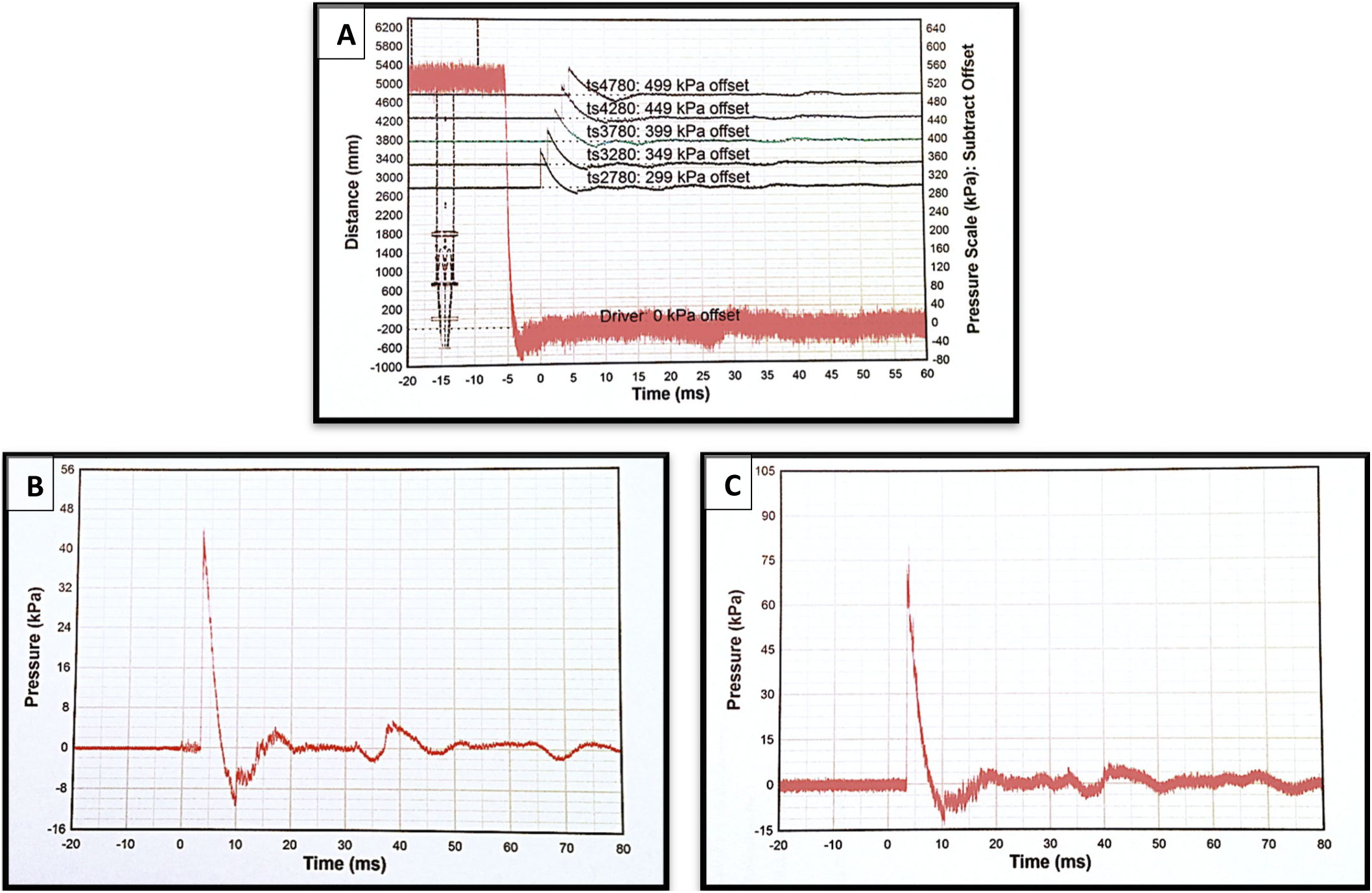
Blast wave characterization. (A) Pressure gauge measurements for a 10 psi blast wave. Pressure sensors are placed along the blast tube, with the lowest pressure recorded by the first sensor closest to the membrane, and the highest pressure recorded by the last sensor, at the end of the tube behind the mouse placement. Sensor ts4280 corresponds to the level of the mouse. (B) Pressure trace, 6 psi. Discarding the 3 highest and 3 lowest of the peaks measured by the sensor closest to the mouse, the blast wave was measured as 41 kPa (6 psi). (C) Pressure trace, 10 psi. Discarding the 3 highest and 3 lowest of the peaks measured by the sensor at the mouse, the blast wave was measured as 69 kPa.

### Acute neurological findings

Mice were initially assessed after blast exposure for their ability to right from a supine position. Tail and corneal reflexes were found to be recovered within seconds of, or simultaneous to, recovery of righting reflex. Time to right after rscBINT was statistically similar to sham, although there were non-significant increases in time to right for the first and second BINTs (Figure 3A). Similarly, smBINT (10 psi) did not increase time to right, indicating that both rscBINT and smBINT produced subconcussive and mild injuries (respectively) without an extended period of unconsciousness at any repetition of BINT.

**Figure 3.**
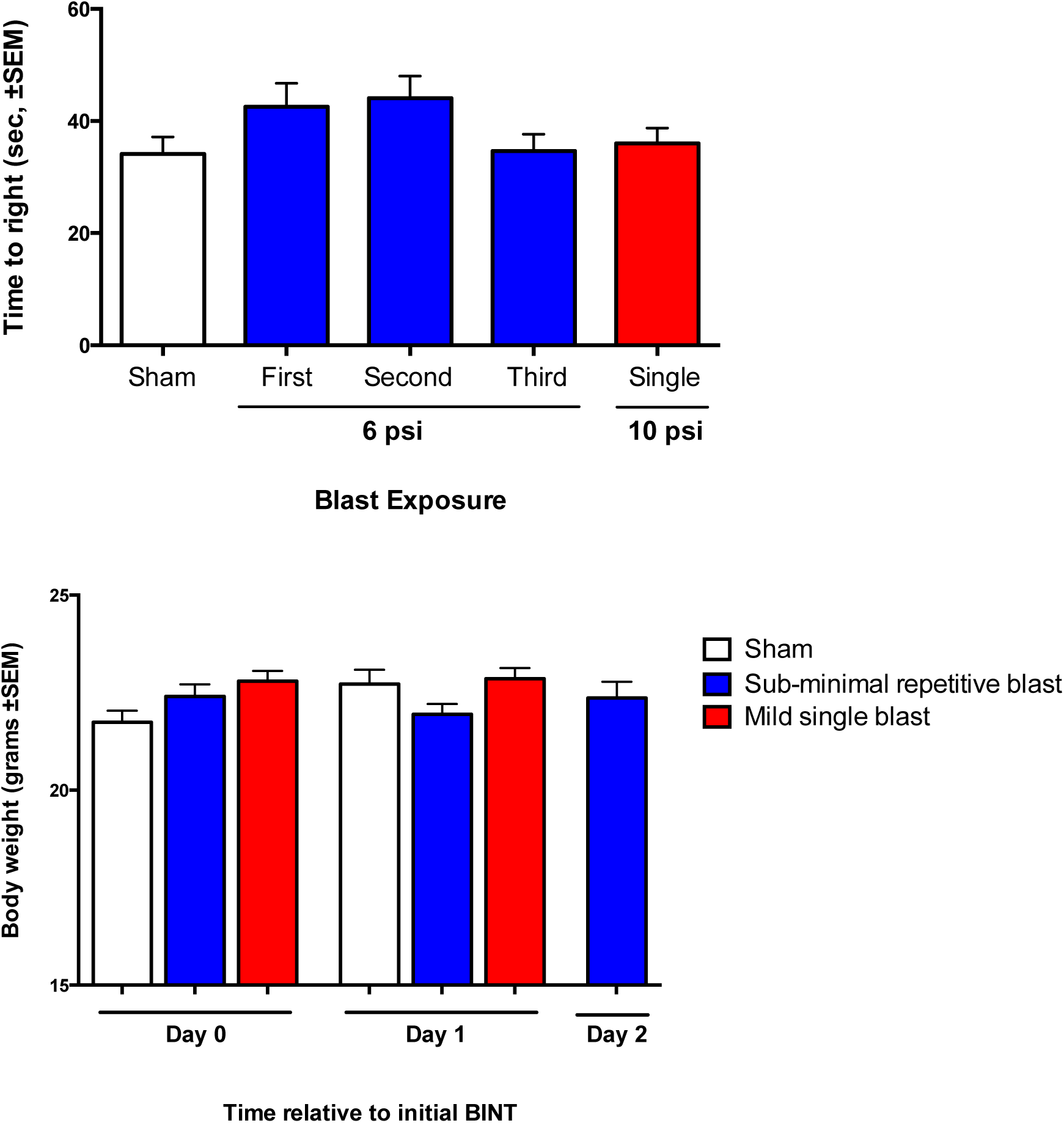
Acute neurological consequences of rsBINT. (A) Time taken from blast exposure to right from a supine position. No significant difference in righting time was detected from sham any stage of the repetitive BINT paradigm. (B) Acute changes in body weight after rsBINT. No significant changes in body weight were observed in mice exposed three rsBINT exposures.

### Acute effects of rscBINT on body weight

Mice were assessed for changes in body weight in the acute period post rscBINT and smBINT (Figure 3B). Sham control mice gained an average of 0.98 g the day following sham BINT induction, and maintained a steady weight increase thereafter. Mice exposed to rscBINT, who had two rscBINT exposures separated by 2 h on the initial day of injury, were found to lose an average of 0.45g between day 0 and day 1 (day of 3^rd^ BINT), however, this loss was not statistically significant. This mild weight loss was also found to be a transient phenomenon, with mice assessed on day 2 (24 h after 3^rd^ exposure) regaining the 0.45g lost the previous day, and gaining weight at a standard pace thereafter. Mice exposed to smBINT maintained a constant weight the day after exposure, before gaining weight again at a standard pace from day 2 onwards.

### Yelverton whole body blast pathology scoring

Examination of whole-body pathology revealed frequent lung injury, with petechial hemorrhages observed 24 hours after rscBINT and smBINT (Figure 4). Pathological sequelae were also detected in the liver and kidneys. When assessed using 2-way ANOVA, a significant effect of both treatment (p<0.0001) and time (p<0.01) were observed, with no interaction between the factors. Tukey’s post hoc analysis determined significant increases in blast-induced pathology in smBINT and rscBINT mice at 24h when compared to sham (p < 0.01 and p < 0.001), with this pathology decreasing when examined at 7d.

**Figure 4.**
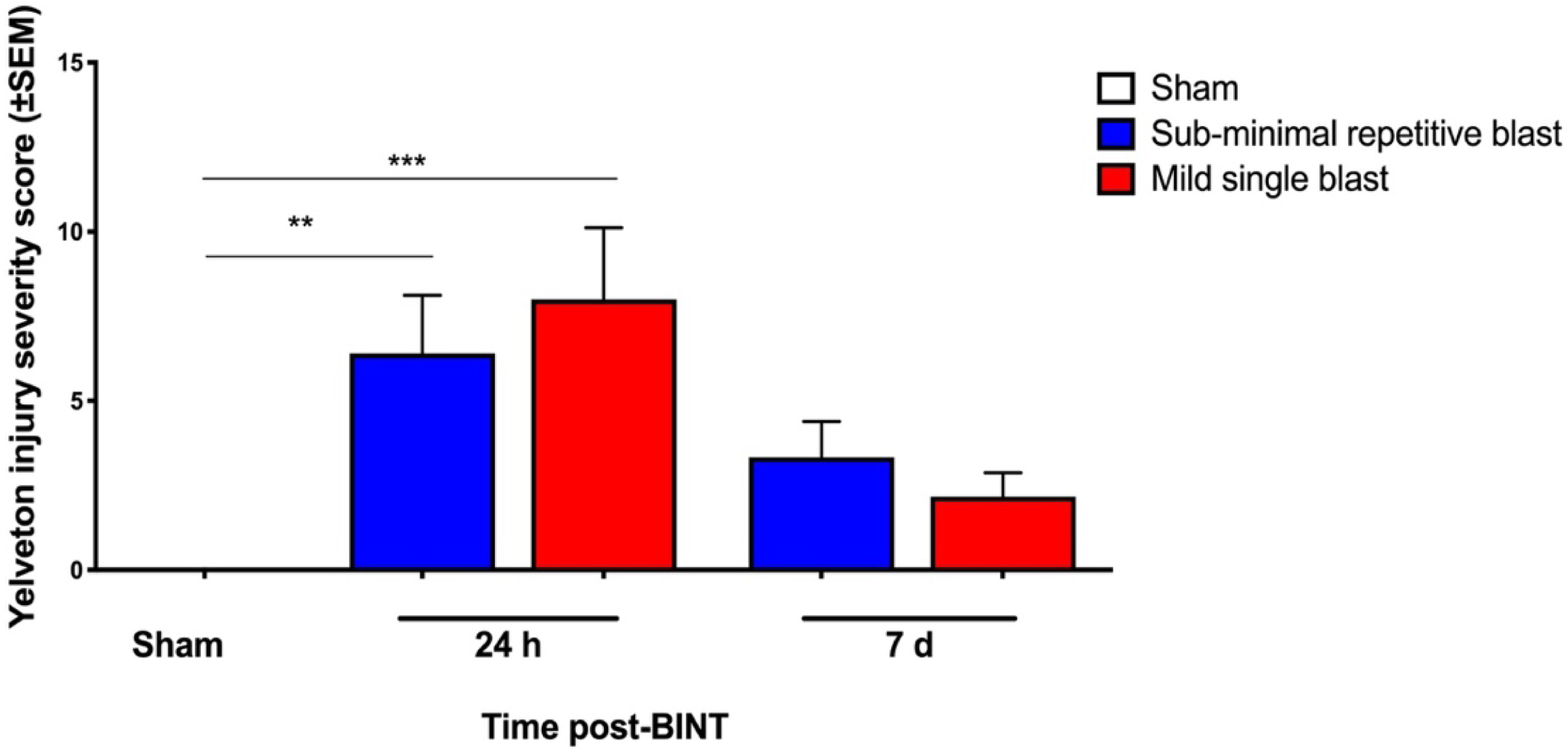
Longitudinal Yelverton injury severity scores for rsBINT and mBINT. Significant effects of both treatment and time on whole-body pathology were detected by 2-way ANOVA. ^**^p < 0.01, ^***^ p < 0.001.

### Effects of rscBINT and smBINT on cerebellar Purkinje neurons

Histological examination of cerebellar tissue with H&E stain revealed alteration in uniformity of Purkinje cells 24h after either rscBINT or smBINT (Figure 5), with a haphazard distribution of Purkinje neurons alongside pyknotic neuronal bodies in the cerebellar lobules. This pathology was also overserved at 7d, with disorganization more prominent, a visible gap between the Purkinje and granular cell layers, and areas of apparent loss of groups of Purkinje neurons.

**Figure 5.**
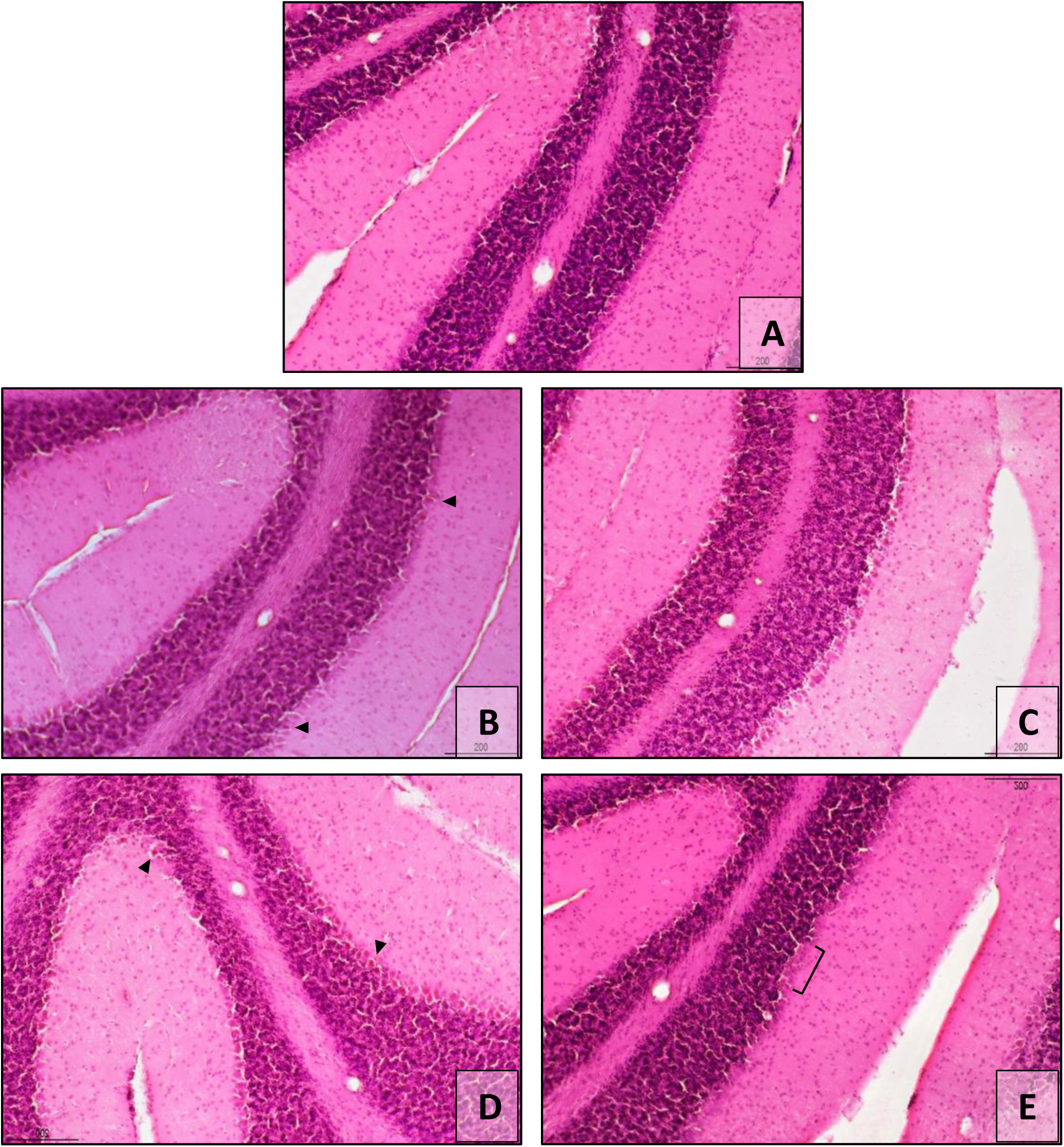
Cerebellar Purkinje cell pathology after rscBINT and smBINT. The Purkinje cell layer appeared uniform with H&E stain in sham (A) mice, while numerous cells appeared pyknotic (arrowheads) with a slightly disorganized distribution 24h after rscBINT (B) or smBINT (D). This pathology persisted at 7d, with increased space between the Purkinje and granular cell layers after rsBINT (C) and visible gaps of missing cells after mBINT (E).

## Discussion

In this paper we describe a new militarily relevant mouse model of BINT, which faithfully reproduces subconcussive and mild injuries. Previous research on BINT has primarily focused on moderate to severe blast injuries, with comparatively little known about the consequences of subconcussive and mild BINT. While characterization of all injury severities is important, models of injuries on the milder end of the spectrum are of vital importance because such injuries occur with higher frequency, and comprise more than 80% of recorded BINT injuries^39^. Blast exposures are frequent in theatre and in the course of training especially among military personnel involved in artillery training, and recent data show that low-level repeated overpressure exposure of soldiers can result in transient symptomatology that overlaps with sub-concussive like effects^54^. The need for preclinical models capable of mimicking real-life scenarios as closely as possible has been underlined during the 2015 event organized by the United States Department of Veterans Affairs (VA) Office of Research and Development (ORD) as well as in the consensus paper summarizing the discussions^55^. Greater emphasis on the effects of repeated, low-level primary blast exposures, such as those related to weapons firing on brain functions, in preclinical model development was one of the key recommendations.

Our model employs a spherical blast wave to which the animal faces head-on in the ‘standing’ bipedal position, rather than the more typically modeled quadrupedal stance. Animal positioning for BINT modeling has been an issue of contention, with most models positioning quadrupedal rodents and larger animals in more naturalistic body positions in which the head first and primarily interacts with the blast wave. In contrast, we believe that positioning quadrupedal animals to approximate bipedal stance better models the clinical scenario, because the effects of the blast wave on the head cannot easily be uncoupled from the effects of the blast wave on the body^56^. Our BINT model also differs from other models by use of a whole-body restraint which provides several important benefits for modeling primary blast: it restrains the animal without the use of materials which may impede or redirect passage of the blast wave through the body; it prevents excessive ‘bobble head’ acceleration movement which confounds primary blast pathology^57-59^; and it does not rely on restraint at the limbs, which would likely increase systemic inflammation and cause localized limb damage, which may then misrepresent the inflammatory state and could be falsely attributed as sensorimotor dysfunction. The structural design of our animal holder also allowed for direct administration of fast-acting controllable anesthetic via nose cone, avoiding the need for long-lasting injectable anesthetics and rapid recovery from the short injury induction.

While parameters for mild BINT are more readily distinguishable from injuries of higher severity, subconcussive BINT is more challenging to model, with no ready criteria for an injury that is (by definition) unobservable ^60^. In order to examine subconcussion, our primary criterion was that a single subconcussive injury would be conducted at the highest pressure for which there was no observable neurological deficit. We found that this correlated to 41 kPa (6 psi). Employed as a repetitive injury, we found that rscBINT did not significantly alter duration of time-to-right after one, two or three rscBINT exposures, with times equivalent to those observed for sham or smBINT. This indicates that our model induced acute neurological dysfunction which was almost unrecognizable immediately after the insults. Likewise, we found no reduction of body weight after a smBINT or any iteration of 1, 2 or 3 rscBINTs, suggesting an absence of severe systemic complications, and resumed characteristic food and water intake as well as rapid recovery comparable to that of sham animals. Taken together, these findings parallel the information soldiers exposed to repetitive low-intensity blasts provide suggesting that this model could be useful to discern subtle cerebral and systemic changes due to repetitive low-intensity blasts.

While neither smBINT or rscBINT induced observable neurological dysfunction, examination of whole-body blast pathology demonstrated peripheral organ pathology similar to that observed after BINT in the military scenario^2,10^, with superficial petechial hemorrhage most frequently in the lungs and (to a lesser extent) the liver and kidneys. Using Yelverton’s pathology scoring system for blast injuries, the severity index score represented mild pathology which was most prominent 24 hours after smBINT or the 3^rd^ rscBINT, and reduced by approximately 50% by 7 days post-injury, indicating a resolution of injury over time. Peripheral injuries are a common feature of whole-body blast, with the lungs particularly susceptible due to blast overpressure and characteristic development of edema and hemorrhage accompanied by inflammatory cell infiltrates^20,61,62^. While lung pathology is progressive in more severe injuries, the superficial nature of injury and rapid pathological resolution observed in this model speaks to the mild nature of injury.

We examined primary pathological change in the Purkinje neurons of the cerebellum based on the particular cerebellar vulnerability to mild blast that has now been shown in several preclinical studies^20,28,63^. We found that three episodes of rscBINT and a single smBINT induced structural alteration of the Purkinje cell layer, with pyknotic neurons visible in both injuries at 24h, and areas of Purkinje loss persisting at 7d. These findings confirm that our model reproduces these findings and add to the growing body of literature which validates findings of cerebellar pathology in military personnel and veterans with blast injuries^11,27,28^. A recent study examining cerebellar tissue from veterans and active military with history of blast demonstrated dystrophic Purkinje cell arbor morphology and loss in expression of the neuronal glutamate transporter EAAT4^63^. Reductions of anisotropy have been found in the middle cerebellar peduncles using diffusion tensor imaging acutely and sub-acutely after mild blast in military populations^64^, with the middle cerebellar peduncle as the primary driver of altered diffusivity, indicative of primary axonal injury in US military personnel 2-4 years post-injury^27^. FDG-PET studies of clinical veterans many years after blast have also described decreased cerebellar glucose metabolism bilaterally in the cerebellum and cerebellar vermis^11^, suggesting that cerebellar pathology may persist chronically after injury.

## Conclusion

This work represents a new model of militarily relevant primary BINT which faithfully reproduces low-level primary blast-induced repetitive and subconcussive injuries without overt neurological dysfunction, but with significant early multi-organ responses. Using readily sourced and customizable materials, this model places mice in a bipedal stance facing into the blast wave, as a more accurate representation of the real-life clinical scenario.

